# Energy Flux Regulates Cell Death Induced by California Serogroup Orthobunyaviruses

**DOI:** 10.64898/2026.03.13.711679

**Authors:** LM Francomacaro, H Matey, KS Beauchemin, AB Evans, S Supattapone

## Abstract

The California serogroup (CSG) of orthobunyaviruses includes neuroinvasive viruses with varying pathogenicity. La Crosse virus (LACV) is a leading cause of pediatric arboviral encephalitis in the USA, while Inkoo virus (INKV) is widespread in Northern Europe but rarely causes disease. The reassortment potential of CSG viruses raises concerns about emerging virulent strains and highlights the need to develop therapies that are broadly effective against multiple CSG viruses. To identify host factors mediating viral neurotoxicity, we performed genome-wide CRISPR-Cas9 knockout screens in human neuroblastoma cells infected with LACV or INKV. Analysis revealed largely overlapping host dependency factors for both viruses. Unexpectedly, the screens identified mitochondrial energy production as a major pathway required for both LACV- and INKV-induced cell death. Reducing host cell energy production with mild hypothermia or sugar source substitution prolonged cell survival during viral infection with additive effects mediated by different mechanisms. Both manipulations also protected neuroblastoma cells from the Bunyamwera virus (BUNV), a non-CSG orthobunyavirus; and mild hypothermia protected mature human neurons from LACV. These results highlight host energy metabolism as a key modulator of CSG virus cytotoxicity and suggest novel avenues for general non-invasive therapeutic intervention against current and future strains of these and other orthobunyaviruses.

## Introduction

The California serogroup (CSG) of orthobunyaviruses is comprised of neuroinvasive viruses with varied pathogenicity in humans. These viruses have a documented ability to reassort both *in vitro* and *in vivo* as well as increasingly overlapping endemic regions due expanding geographic ranges for vector species and abundant global travel [1, 6, 11, 31, 42, 46, 49, 60, 66, 84, 86, 116]. The combination of reassortment potential and endemic spread threatens the emergence of novel virulent strains.

La Crosse virus (LACV) and Inkoo virus (INKV) are both CSG viruses with markedly different propensities for causing clinical disease. LACV is a leading cause of pediatric arboviral encephalitis (LACV-E) in the United States, while INKV, though highly seroprevalent in Northern Europe, rarely causes illness requiring medical intervention [13, 20, 25, 39, 70, 77, 78, 83, 92, 106]. These differences in human disease cases are recapitulated in mouse and human neuronal cell models of infection, where LACV is significantly more neurotoxic than INKV [27, 28].

A handful of studies have tested potential antivirals but supportive measures for infected patients remain the clinical standard of care, and no vaccines exist; therefore, avoiding vector species is the primary method of prevention [16, 32, 71, 75, 76, 90]. Perhaps the closest the field has come to a treatment was a clinical trial for ribavirin, which had to be discontinued due to the adverse effects seen at a therapeutic dose [16, 71]. Some studies have found promising pharmaceutical agents by repurposing known antivirals, such as influenza endonuclease inhibitors that impair the cap-snatching required for viral transcription or molnupiravir which causes lethal viral mutagenesis [32, 75]. Other studies have screened potential drugs to identify those that limit LACV cytotoxicity in cell culture and animal models, including the potassium ionophore valinomycin and the Golgi traffic impairing drug rottlerin [76, 90]. One study attenuated the neuroinvasive capability of LACV by targeting mutations to the viral Gc fusion peptide[47]. Despite these strides in identifying potential therapeutics, the standard of care remains supportive.

Since most clinically severe LACV-E cases occur in rural areas and frequently result in long-term neurological sequelae, there is a pressing need for effective and accessible medical interventions [14, 45, 65, 97]. A better understanding of how LACV and INKV interact with host cell machinery will inform not only potential treatments for LACV-E, but also potential treatments for other orthobunyaviruses and any reassortant viruses that may arise.

Several small-scale host genetic studies have examined how CSG viruses interact with host cell machinery. Early work identified apoptosis as a major mechanism of virally-associated cell death and found a protective role for the expression of anti-apoptotic bcl-2 [80]. A targeted look at the innate immune signaling pathways found roles for RIG-1 (a pattern recognition receptor), MAVS (an antiviral signaling protein), and SARM1 (an adaptor protein) in neuronal cell death upon LACV infection [73]. A transcriptomics and targeted siRNA screen found CX43 (a gap function connexin) expression protective in weanling mice and loss of Efna2 (an Ephrin signaling molecule) deleterious in adult mice in the context of LACV infection [8]. A more general look at the total oxidative damage burden in the host cell found that differentiation of neuroblastoma cells via retinoic acid treatment decreased both oxidative damage and susceptibility to LACV induced cell death [81].

No whole genome host cell genetic studies have been conducted specifically for CSG viruses. A couple of such screens have been conducted for other viruses and their findings extrapolated to LACV infection. A screen for Rift Valley Fever phlebovirus, a distantly related bunyavirus, found that knockout of *WDR7* (WD repeat protein family gene potentially involved in endocytosis or secretion) led to impaired LACV viral replication in A549 cells [7]. Likewise, a screen with Schmallenberg virus, a more closely related orthobunyavirus, found that depletion of heparin sulfate at the cell surface downstream of PAPST1 (a sulfotransferase) action decreased LACV infection in HEK293T cells [103]. No studies have examined any Bunyavirus infection in whole genome knockout screens of neuronal cells.

To identify host cell machinery involved in CSG-induced cell death, we utilized whole genome CRISPR-Cas9 knockout survival screens in neuroblastoma cells for each LACV and INKV, comparatively. Although CSG viruses are known to infect several cell types, including muscle cells at the site of mosquito bite, they preferentially infect and kill neurons, leading to clinical encephalitis. The CSG viruses are restricted to replication in neurons in the CNS *in vivo,* and previous studies have shown that human neuroblastoma cells recapitulate the differences in virulence between LACV and INKV. Therefore, we chose human neuroblastoma cells for these screens to identify host factors that mediate neuronal death to a highly neurovirulent (LACV) compared to a mildly neurovirulent (INKV) viruses [9, 18, 28, 91, 112]. To our knowledge, the genetic screens reported herein represent the first whole genome host cell survival screens conducted for CSG viruses.

## Materials and Methods

### Cell Culture Conditions

BE(2)- C (ATCC, catalog #CRL-2268) human neuroblastoma cells were cultured in a 1:1 mix of EMEM (ATCC, catalog #30-2003) and F-12 (Gibco, catalog #11-765-054) media supplemented with 10% fetal bovine serum (Cytiva, catalog #SH30910.03) at 37°C with 5% CO_2_ until >90% confluent then split 1:4-1:10 with 0.05% Trypsin / 0.53 mM EDTA (Corning, catalog #25052CV).

Vero cells (gift from Alyssa Evans, Montana State University) were cultured in EMEM media supplemented with 10% fetal bovine serum (FBS) at 37°C with 5% CO_2_ until >90% confluent then split 1:5-1:20 with 0.05% Trypsin / 0.53 mM EDTA.

WTC-11 NGN2 induced pluripotent stem cells were a gift from Li Gan (Weill Cornell, NY). Cells were grown on plates coated with hESC-qualified Matrigel (Corning, catalog #354277) in mTeSR1 media (StemCell Technologies, catalog #85857), and split with ReLeSR (StemCell Technologies, catalog #100-0483) in the presence of Y-27632 ROCK inhibitor (StemCell Technologies, catalog #72302) per manufacturer’s recommended protocols.

Both BE(2)-C neuroblastoma and WTC11-NGN2 induced neurons (iNeurons) were highly susceptible to infection by LACV and INKV; none of the cell culture treatments caused a change in BE(2)-C morphology (Supplemental Figure 1A-H).

### iPSC differentiation via Doxycycline to iNeurons

WTC11-NGN2 iPSCs were differentiated into mature glutamatergic cortical neurons via doxycycline inducible neurogenin 2 (NGN2) as described by Fernandopulle et al. [33]. Briefly, iPS cells were lifted with Accutase (StemCell Technologies, catalog #07922) with DNase (Sigma, catalog #11284932001) to a single cell suspension and re-plated on Matrigel coated plates in induction media consisting of DMEM/F-12 (Gibco, catalog #11330032) with 1x N2 supplement (Gibco, catalog #17502048), 1x non-essential amino acids (Gibco, catalog #11140050), and 1x L-glutamine (Gibco, catalog #25030081). Induction media was refreshed daily for 3 days with 2 ug/mL doxycycline (Sigma, catalog #D9891) and 10 µM ROCK inhibitor only for day one. Nascent iNeurons were lifted again with Accutase plus DNase and seeded on plates double coated with poly-L-ornithine (Sigma P3655) and laminin (Gibco, catalog #23017015) in cortical neuron media consisting of BrainPhys media (StemCell Technologies, catalog #05790) with 10 ng/mL BDNF (PeproTech, catalog #450-02), 10 ng/mL NT-3 (PeproTech, catalog #450-03), 1 ug/mL laminin, and 1x B27 supplement 17504044). Half-volume media changes were conducted twice weekly; cells were mature at 14 days post-induction start.

### Viral Stocks

LACV (human 1978 strain) and INKV (strain SW AR 83-161) were kindly provided by Alyssa Evans (Montana State University) and Karin Peterson (Rocky Mountain Laboratories NIAID). BUNV (strain 6547-8) was kindly provided by the World Reference Center for Emerging Viruses and Arboviruses (University of Texas). Viral stocks were stored at −80°C, thawed at 37°C prior to inoculations, and discarded after a single freeze/thaw.

### Viral Amplification

Vero cells (<20 passages) were plated to >90% confluence in media with reduced serum (2% FBS) and inoculated at multiplicity of infection (MOI) = 0.1 to MOI = 0.5 with LACV, INKV, or BUNV then incubated at 37C, 5% CO_2_ until cytopathic effect via microscopic visualization was greater than 80%. Media was collected, spun at 500xg for 7 minutes to remove cellular debris, flash frozen via liquid nitrogen submersion, and stored at −80°C to be titered prior to use.

### Viral Plaque Titer

24 well plates were seeded with low passage Vero cells at 100% confluence in reduced serum (2% FBS) media then inoculated in triplicate with a dilution series of virus. A one-hour incubation with intermittent agitation to allow for viral adherence was followed by 2.5x volume addition of 1.5% weight/volume carboxymethyl cellulose (Sigma, catalog #C4888) in minimal essential media (Gibco, catalog #12360038). Plates were incubated at 37°C, 5% CO_2_ without disruption for 2-3 days prior to fixation with 7.5% formaldehyde for 1 hour at room temperature and plaque visualization with 0.35% weight/volume crystal violet in 95% ethanol. Viral titers were calculated from replicate plates in plaque forming units (PFU) per mL.

### Cas9 Clonal Cell Line Creation and Functional Validation

BE(2)-C cells were engineered to constitutively express Cas9 via lentiviral transduction of lentiCRISPRv2 blast (Addgene, catalog #98293), which was a gift from Brett Stringer [96]. A clonal line of BE(2)-C Cas9 cells was generated via single cell outgrowth. Function of the Cas9 enzyme in the clonal line was confirmed using the Cellecta transduction and flow cytometry based CRISPRuTest (Cellecta, catalog #CRUTEST) (Supplemental Figure 2A-B).

### Whole Genome Knockout Library Creation

Human Brunello CRISPR knockout pooled library was a gift from David Root and John Doench (Addgene, catalog #73178) [22]. The Brunello plasmid was amplified in Endura electrocompetent cells (Lucigen, catalog #60242-2) and harvested with a Plasmid Maxiprep Kit (Qiagen, catalog #12162) prior to third generation lentiviral packaging in HEK239FT cells (ThermoFisher, catalog #R70007). A functional titer in BE(2)-C Cas9 cells was performed to achieve an MOI = 0.3 with the Brunello library lentivirus. The library was cultured for several passages prior to genomic DNA extraction with a Blood and Cell Culture midi kit (Qiagen, catalog #13323); the gDNA was subjected to PCR preparation and next generation sequencing as detailed below to confirm balanced representation of single-guide RNAs (sgRNA). One copy of the library was defined as 500-fold representation per sgRNA: The Brunello library consists of 76,441 sgRNAs; a single copy of the Brunello library was defined as 38,220,500 cells, rounded to 40 million cells for ease. During culture, passage, and cryopreservation, at least one library copy with 500-fold representation was maintained to protect representation.

### Targeted Membrane Protein Knockdown Library Creation

BE(2)-C cells were engineered to constitutively express dCas9-KRAB via lentiviral transduction of pLX_311-KRAB-dCas9 (Addgene, catalog #96918), which was a gift from John Doench and William Hahn and David Root [88]. A clonal line of BE(2)-C dCas9-KRAB cells was generated via single cell outgrowth. Function of the Cas9 enzyme in the clonal line was confirmed using the Cellecta CRISPRiTest (Cellecta, catalog #CRITEST).

Human subpooled CRSIRPi-V2 membrane proteins library was a gift from Jonathon Weissman (Addgene, catalog #83976) [48]. The membrane protein plasmid library was amplified in Endura electrocompetent cells (Lucigen, catalog #60242-2) and harvested with a Plasmid Maxiprep Kit (Qiagen, catalog #12162) prior to third generation lentiviral packaging in HEK239FT cells (ThermoFisher, catalog #R70007). A functional titer in BE(2)-C dCas9-KRAB cells was performed to achieve a MOI = 0.3. Genomic DNA extraction was performed with a Blood and Cell Culture midi kit (Qiagen, catalog #13323); the gDNA was subjected to next generation sequencing to confirm balanced representation of single-guide RNAs (sgRNA).

### PCR Preparation of gDNA for Illumina Sequencing

Reactions were performed in an environment and using tools sterilized with 10% bleach for 20 minutes then rinsed with molecular biology grade water to remove any trace contamination from prior genomic libraries. Q5 Hot Start High Fidelity 2x Master Mix (NEB, catalog #M0494) was used per manufacturer’s specifications with a reaction volume of 50 μL. Three successive rounds of PCR were used to prepare gDNA for sequencing (Supplemental Figure 3A-D); primers are listed in Supplemental Table 2. Each round of PCR was followed by pooling reaction products and purification with a QIAquick PCR purification kit (Qiagen, catalog #28104). PCR1 extracted sgRNA sequences from genomic DNA using common flanking sequences. PCR1 reactions were performed to consume all gDNA produced by the Qiagen Blood and Cell Culture midi kit, with 1.75 μg gDNA per reaction. PCR2 attached Illumina sequencing adaptors and barcodes. PCR2 reactions, 10 per copy of library, were performed using 10 μL of pooled and purified PCR1 product as input. Uniquely barcoded PCR2 forward primers identified samples, allowing a pooled mix of all 12 PCR2 reverse primers. PCR3 ensured blunt ends to improve sequencing clustering and quality; 16 PCR3 reactions were performed per copy of the library using 150 ng of pooled and cleaned PCR2 product per reaction as input. PCR3 product was pooled and run on a 2% agarose gel prior to gel extraction using a QIAquick Gel Extraction Kit from (Qiagen, catalog #28704) and subsequent concentration of extracted product using a QIAquick PCR purification kit. Resulting samples were ready for sequencing.

### Illumina Next Generation Sequencing

Fragment analyzer and Qubit were run as quality control measures prior to analysis. Sequencing was performed by Dartmouth Genomics Shared Resource Core facilities on an Illumina MiniSeq using high output 75 or 150 cycle run with single end reads. Read count was a minimum of 50 reads per sgRNA or 3.8 million reads per library copy; 10% Phi-X Control (Illumina, catalog #FC-110-3001) was added to increase diversity. Samples were deconvoluted to FASTQ files for subsequent analysis.

### MAGeCK Pipeline Analysis of NGS

Deconvoluted FASTQ files were analyzed using the MAGeCK-VISPR and MAGeCKFlute pipelines [63, 110]. Samples were analyzed as paired replicates; defaults used for all other parameters. A false discovery rate (FDR) < 0.05 was used as a threshold for significant hits.

### LACV Whole Genome Knockout Survival Screen Design

BE(2)-C Cas9 Brunello library cells were passaged with at least one full copy of the library (500x all sgRNA, 39 million cells) to maintain representation. Screens were conducted in biological triplicate as pairs of infected and uninfected samples. Infected replicate one and control replicate one were performed both temporally separate and from a different thawed cell aliquot than infected replicates two and three and control replicate two, accordingly, the samples were analyzed in a paired manner. Cells were plated at about 30-40% confluence (4-5 million cells/15 cm plate) on day −1 with one copy of library per condition and allowed to adhere overnight. On day 0, two plates were counted to estimate cell number per plate to calculate viral inoculation at MOI = 1. The lifted and counted cells were discarded. All plates, infected and control, were refreshed with complete media. Infected plates were inoculated with LACV at MOI = 1. Plates were incubated for 72 hours until about 10% of cells remained viable on the infected plates. Genomic DNA extraction was performed with a Qiagen Blood and Cell Culture midi kit.

### INKV Whole Genome Knockout Survival Screen Design

Performed as described for LACV Survival Screen design with the following modifications. Infected replicates one through three and control replicates one through three were performed at the same time but paired such that numbered samples came from the same thawed cell populations. Post-inoculation, plates were incubated for 7 days until about 10% of cells remained viable on the infected plates. Infected plates were refreshed with a half media spike in on days two and four; no media was removed. Control plates were lifted on days two and four, mixed well, and a representative copy (40 million cells) re-plated. Following incubation, genomic DNA extraction was performed with a Blood and Cell Culture midi kit.

### LACV Targeted Membrane Protein Knockdown Survival Screen Design

BE(2)-C dCas9-KRAB membrane protein library cells were passaged with at least one full copy of the library (500x all sgRNA) to maintain representation. The screen was conducted in unpaired triplicate. Cells were plated at about 30-40% confluence (4-5 million cells/15 cm plate) on day −1 with one copy of library per replicate and allowed to adhere overnight. On day 0, infected plates were inoculated with LACV at MOI = 1. Plates were incubated for 72 hours prior to genomic extraction and sgRNA analysis as described above.

### Cell Counting

All cell counts were performed using a Countess 3 FL Automated Cell Counter with trypan blue dye and disposable chamber slides per manufacturer’s specifications (ThermoFisher, catalog #A49866). Every sample was counted twice and reported as an average of those two counts. Low count samples were spin concentrated prior to counting to ensure accuracy.

### Doubling Time Determination

BE(2)-C cells grown under varied conditions were seeded in well plates at very sparse confluence. Daily counts were taken from triplicate wells and fitted to an exponential y=ae^bx^ from which doubling time was calculated.

### Viral Susceptibility Studies

On day −1, 6 or 12 well plates were seeded with low passage BE(2)-C wild type cells at about 30% confluence and incubated at 37°C, 5% CO_2_ overnight to adhere. On day 0, three representative wells were counted for each condition to allow for MOI calculations, all wells were refreshed, and inoculations were performed. Cells were incubated at 37°C, 5% CO2 unless otherwise noted for 48-96 hours prior to cell counts. Cell survivals were calculated based on treatment matched MOI = 0 controls and compared to wild type survival rates. Cells were counted as described above with trypan blue staining to count only live cells. Experiments were performed in triplicate.

### Viral Susceptibility Studies: Temperature

BE(2)-C wild type cells were maintained at 37°C until day 0 following inoculation. For each 33°C and 37°C, wells were inoculated at the following in triplicate: LACV MOI = 0, 0.0001, 0.01, 1; INKV MOI = 0, 0.01, 1, 100; TAHV MOI = 0, 0.01, 1; BUNV MOI = 0, 0.01, 1. Plates were then incubated at either 37°C, 5% CO_2_ or 33°C, 5% CO_2_ for approximately 48 hours (LACV, TAHV), 72 hours (BUNV), or 96 hours (INKV). Timepoints were selected to capture differential survival percentages for each virus.

WTC11-NGN2 induced neurons (iNeurons) were grown up on 10-cm plates at a low confluence until mature (day 14). They were inoculated with LACV MOI = 10 and incubated at either 37 or 33°C, 5% CO_2_ for approximately 96 hours. Cell survival percentages were calculated based on treatment matched MOI = 0 controls and compared to wild type survival rates. Cells were counted as described above with trypan blue staining to count only live cells. Experiments were performed in triplicate.

### Viral Susceptibility Studies: Carbon Source

10 mM D-(+)-galactose (Sigma, catalog #G0750) or 10 mM D-(+)-glucose (Sigma, catalog #G8270) was added to glucose free DMEM (Gibco, catalog #11966025) and 0.22 μm filter sterilized prior to the addition of 10% FBS to create galactose or glucose media, respectively. BE(2)-C wild type cells were cultured in either glucose or galactose media for a minimum of 3 days prior to studies. Inoculations were at the following in triplicate: LACV MOI = 0, 0.01, 1; INKV MOI = 0, 0.01, 1, 100; TAHV MOI = 0, 0.01, 1; BUNV MOI = 0, 0.01, 1. Plates were then incubated at 37°C, 5% CO_2_ for approximately 48 hours (LACV, TAHV, BUNV), or 96 hours (INKV).

### Viral Susceptibility Studies: Oligomycin

Oligomycin (Fisher Scientific, catalog #J61898MA) was dissolved in dimethyl sulfoxide (DMSO, Sigma, catalog #D2650) to create a 50 mM stock solution. Oligomycin stock or pure DMSO was diluted 1:400 in complete media to create working solutions. At the time of viral inoculation at MOI = 1 (LACV) or 100 (INKV), oligomycin and DMSO working solutions were spiked into wells in triplicate to achieve oligomycin concentrations of 0 or 5 μM balanced for total DMSO dose as well as an untreated wild-type control. The same was done for matched plates receiving no viral inoculations. Plates were incubated at 37°C, 5% CO2 for 48 hours (LACV) or 72 hours (INKV).

### Viral Susceptibility Studies: Caspase 9 Inhibitor

Caspase 9 inhibitor Z-LEHD-FMK (R&D Systems, catalog #FMK008) was dissolved in DMSO to create a 20 mM stock solution. BE(2)-C wild type cells were subjected to conditions of 10 μM Z-LEHD-FMK, DMSO, or untreated, initiated at the time of viral inoculation (LACV MOI = 1). Cells were incubated at 37 °C, 5% CO_2_ for 48 hours, in triplicate.

### Viral Susceptibility Studies: Pan Caspase Inhibitor

Pan Caspase inhibitor Q-VD-Oph (SelleckChem, catalog #S7311) was dissolved in DMSO to create a 50 mM stock solution. BE(2)-C wild type cells were subjected to conditions of 10 μM Q-VD-Oph, DMSO, or untreated, initiated at the time of viral inoculation (LACV MOI = 0, 1). Cells were incubated at 37°C, 5% CO_2_ for 48 hours, in triplicate.

### Viral Susceptibility Studies: Combination of Carbon Source and Temperature

BE(2)-C wild type cells were grown in either galactose or glucose media as described above for at least 3 days. For each glucose and galactose, two plates were inoculated in triplicate with LACV MOI = 0, 0.0001, 0.01, 1. One of each media condition was placed at 33°C, 5% CO_2_, the other at 37°C, 5% CO_2_ for 48 hours.

### Viral Susceptibility Studies: Combination of Pan Caspase Inhibitor and Temperature

Four plates of BE(2)-C wild type cells were inoculated with LACV MOI = 0, 1, 5 in triplicate. Two of the plates were treated with 10 μM Q-VD-Oph pan caspase inhibitor at the time of viral inoculation. One each of Q-VD-Oph treated and untreated plates were placed at 33°C, similarly one each was placed at 37°C, 5% CO_2_ for 48 hours. After incubation, percents survival were determined.

### Viral Susceptibility Studies: Combination of Pan Caspase Inhibitor and Carbon Source

BE(2)-C wild type cells were grown in either galactose or glucose media as described above for at least 3 days. For each glucose and galactose, two plates were inoculated in triplicate with LACV MOI = 0, 1, 5. One plate for each media was also treated with 10 μM Q-VD-Oph pan caspase inhibitor at the time of viral inoculation. Plates were incubated at 37°C, 5% CO_2_ for 48 hours and percents survival were determined.

### Caspase 9 Flow Cytometry Assay

BE(2)-C wild type cells were plated at about 40% confluence on 10-cm plates and allowed to adhere overnight. All plates were refreshed and treated. Treatments were balanced by DMSO dose and include combinations of 4 μg/mL camptothecin (Fisher Scientific, catalog #J62523MD), 10 μM Q-VD-Oph, LACV MOI = 1, 5 μM oligomycin. Additionally, some BE(2)-C cells were pre-cultured in galactose media for 3 days prior. Some BE(2)-C cells were incubated at 33°C rather than the standard 37°C. Approximately 24 hours after treatment initiation, the Green Fluorescent FAM-FLICA Caspase-9 (LEHD) Assay kit was used according to manufacturer’s instructions (Antibodiesinc, catalog #913). An unstained DMSO condition was also included. Stained and fixed samples were analyzed on a CytoFLEX LX (Beckman Coulter) flow cytometer alongside AccuCheck ERF Reference Particles (ThermoFisher, catalog #A55950) with laser settings FSC 50, SSC 150, and B525 FITC 30. Resulting data was analyzed using FlowJo version 10.10.0 to gate for single cells and determine median fluorescence intensities.

### Reduction Potential Studies

BE(2)-C wild type cells were pre-grown in typical culture conditions, 33°C incubation, galactose media, or the combination of 33°C incubation and galactose media. Cells were seeded in 24 well plates, allowed to adhere, and grown for 2 days. PrestoBlue HS Cell Viability Reagent (ThermoFisher, catalog #P50200), a resazurin live cell reduction potential dye, was warmed to room temperature and added at 1/10 culture volume. The dye was incubated, protected from light, at 37°C or 33°C for 2.5 hours. Plates were read in a Molecular Devices Spectra Max iD5 plate reader using the monochromator set to 560/590 nm. Fluorescence values were adjusted based on average fluorescence from media only wells and scaled to cell count prior to inter-sample comparison.

### Mitochondrial Membrane Potential Studies

BE(2)-C wild type cells were pre-grown in typical culture conditions, 33°C incubation, or galactose media. Cells were seeded in 24 well plates, allowed to adhere, and grown for 2 days. A JC-10 based mitochondrial membrane potential kit (Sigma, catalog #MAK159) was used per manufacturer’s instructions. Briefly, aggregates of JC-10 in intact mitochondria are red fluorescent while monomeric JC-10 from leaky mitochondria is green fluorescent. 100x JC-10 was diluted in Assay Buffer A, then added to wells at ½ culture volume and incubated, protected from light, at 37°C or 33°C for 30-60 minutes. Assay Buffer B was added 1:1 to Assay Buffer A. Fluorescence was read in a Molecular Devices Spectra Max iD5 plate reader using the monochromator set to 490/525 nm and 540/590 nm.

### Statistical Analyses

Tests of significance were performed in Prism version 10.4.1 as noted in the results section.

## Data Availability

The raw sequencing data for both screens are available as Mendeley deposits under the following DOIs:

Control library: 10.17632/m8pkzbktbk.1
LACV Screen: 10.17632/3n5hzxt58t.1
INKV Screen: 10.17632/8cj9tzb4pz.1

## Acknowledgements

The authors thank Karin Peterson (Rocky Mountain Laboratories, Hamilton, MT) for helpful discussions. This study was funded by the National Institute for Neurological Diseases and Stroke (1R37NS125431, R01NS117276 and R01NS118796 to S.S.) and the National Institutes of Health (P20-GM113132 to Dean Madden)**. Additionally, NGS** was carried out in the Genomics and Molecular Biology Shared Resource (RRID:SCR_021293) at Dartmouth which is supported by NCI Cancer Center Support Grant 5P30CA023108 and NIH S10 (1S10OD030242) awards.

The lentiCRISPRv2 blast plasmid (Addgene, catalog #98293) was a gift from Brett Stringer. The human Brunello CRISPR knockout pooled library was a gift from David Root and John Doench (Addgene, catalog #73178).

None of the authors have conflicts of interests.

## Results

### Genome-wide knockout survival screen of LACV-infected cells identified pro-apoptotic and mitochondrial energy production pathways as major effectors of viral pathology

To identify genes associated with susceptibility to California serogroup orthobunyaviruses, we performed a genome-wide CRISPR knockout survival screen on LACV-infected neuroblastoma cells. The Brunello whole genome knockout library was inserted into a monoclonal BE(2)-C Cas9 line via low titer lentiviral transduction; these cells are henceforth referred to as library cells [22]. To allow for enrichment of genes associated with viral susceptibility, library cells were infected with LACV MOI = 1 in paired triplicate and allowed to succumb to infection for 72 hours (Figure 1). Post-infection incubation of 72 hours resulted in about 10% survival for LACV infected library cells; further incubation resulted in total loss of all library cells (Supplemental Table 1; Supplemental Figure 1A-C; Supplemental Figure 2C). Thus, single gene knockouts were not sufficient to prevent viral cell death.

**Figure 1.**
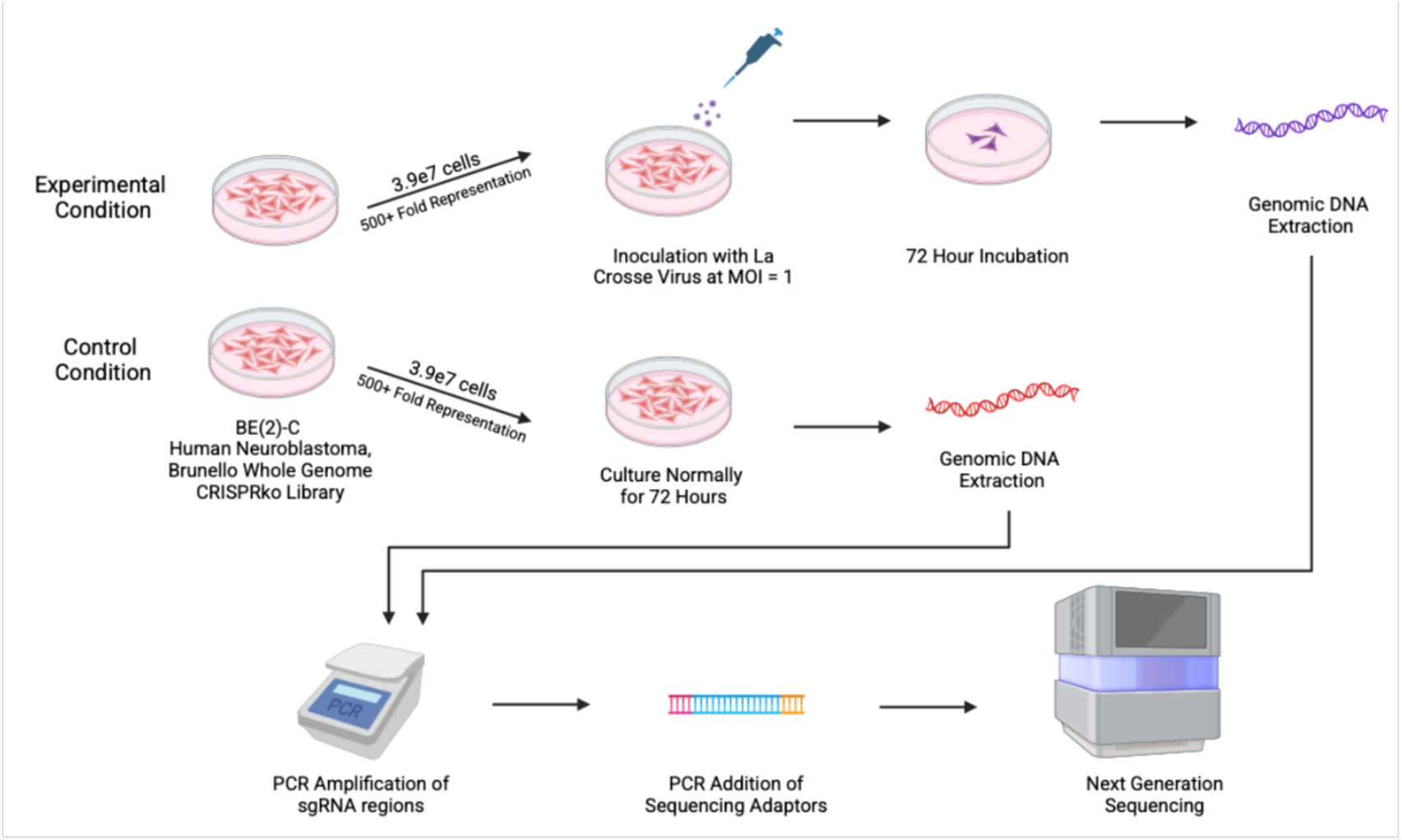
Schematic of LACV Survival Screen Design. Both experimental and control conditions were conducted in paired triplicate. The INKV survival screen differed from the LACV outlined above only for incubation period, which was 7 days rather than 3 days. Details of PCR amplification are in Supplemental Figure 3

Raw sequencing data for the unselected library and three screen replicates are deposited with Mendeley [35, 37] DOI: 10.17632/m8pkzbktbk.1 & 10.17632/3n5hzxt58t.1. Analysis of NGS results using the MAGeCK-VISPR pipeline identified 153 genes for LACV that met the significance threshold of FDR < 0.05; these genes are referred to as hits (Supplemental Table 3). Since cells with hit genes knocked-out survived viral infection for longer than either wild type cells or cells with different knocked-out genes, the genetic hits are associated with increased susceptibility to viral infection when intact.

The genetic hits were ranked by statistical significance, with higher ranked hits showing a stronger, more consistent enrichment across replicates and same gene targeting sgRNAs. Strong hits had nearly universal enrichment of the four unique sgRNAs per gene across all three replicates (Figure 2A-B). Pathway analysis conducted via MAGeCK-VISPR using the KEGG, GOBP, REACTOME, and Complex databases revealed robust action through apoptotic and metabolic pathways, particularly caspase 9-dependent apoptosis and mitochondrial energy production (Figure 2C) [52–54]. Thus, library cells with impaired apoptosis or energy production machinery survived LACV infection longer than wild type or other library cells.

**Figure 2.**
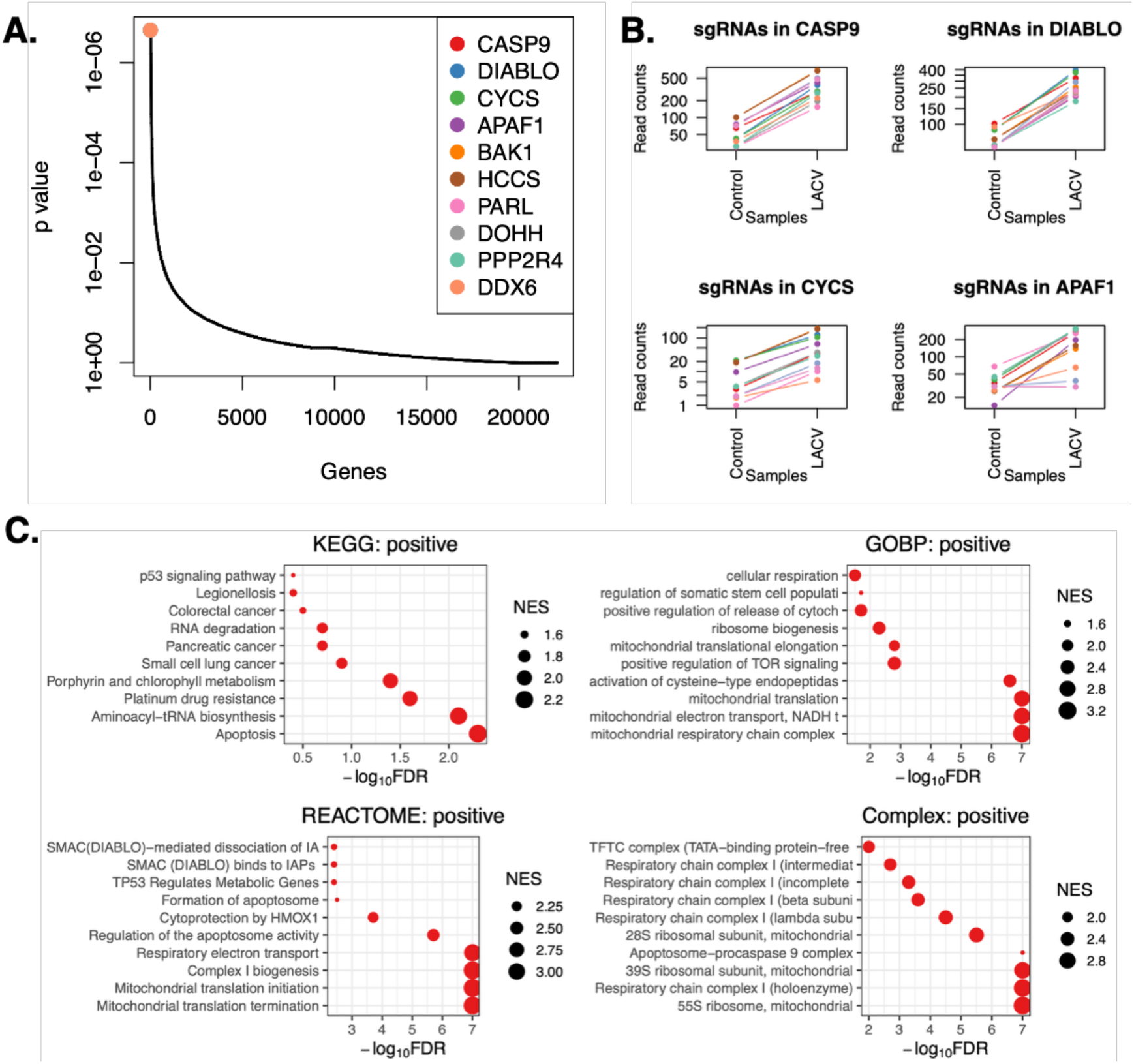
LACV Survival Screen Genetic and Pathway Hits. (A) MAGeCK-VISPR analysis of NGS results from the LACV whole genome knockout survival screen were plotted as a distribution of *p* values across genes, with the top ten hits highlighted. (B) Read count plots from the top four ranked hits from the LACV Screen show nearly universal enrichment from control to LACV samples; each line represents one sgRNA from one replicate. (C) Pathway analysis of genetic hits was conducted using the Kyoto Encyclopedia of Genes and Genomes (KEGG), Gene Ontology Biological Process (GOBP), Reactome, and Complex databases through the MAGeCK-VISPR platform. Negative log_10_ false discovery rates (FDR) were plotted for pathway hits with point size corresponding to relative normalized enrichment score (NES).

### Screen hits shared between LACV and INKV revealed a common cytotoxic mechanism

Given their disparate propensities for causing human disease that rises to the level of clinical significance and documented differences in virulence in mice and human neuronal cells, we investigated whether LACV and INKV utilized different pathways for inducing cell death. INKV MOI = 1 required a longer incubation with library cells to reduce cell populations to 10% survival, 7 days compared to 3 days for LACV (Supplemental Figure 2C). Of note, wild type cells cultured at the same conditions and timescale of the INKV 7-day infection span do not die due to over-culturing. Despite a slower cytotoxic effect, INKV also killed all library cells if given sufficient incubation time: demonstrating once again that single gene knockouts were not sufficient to prevent viral cell death entirely.

Raw sequencing data for the unselected library and three screen replicates are deposited with Mendeley [35, 36] DOI: 10.17632/m8pkzbktbk.1 & 10.17632/8cj9tzb4pz.1. The INKV survival screen had 303 genes that met the significance threshold of FDR < 0.05, nearly double the number of hits found in the LACV screen (Supplemental Table 4). Of the top 10 hits from the INKV screen, 8 are shared with the top 10 hits from the LACV screen (Figures 2A, 3A). Of all hits, 89 are shared between viruses (Supplemental Tables 3&4). INKV top hits showed similarly strong enrichment across replicates and same gene targeting sgRNAs (Figure 3B). Strikingly, the INKV screen identified many identical pathways as the LACV screen (Figures 2C, 3C). There were no remarkable single gene or pathway hits that differed between the two viruses. Of note, top ranked pathways that may appear to be unique, such as “positive regulation of TOR signaling” for LACV are shared as statistically significant hits, but simply ranked differently for INKV (Figures 2C, 3C). In combination, the two survival screens discovered shared mechanisms of cytotoxicity, mainly apoptosis and respiratory energy production.

**Figure 3.**
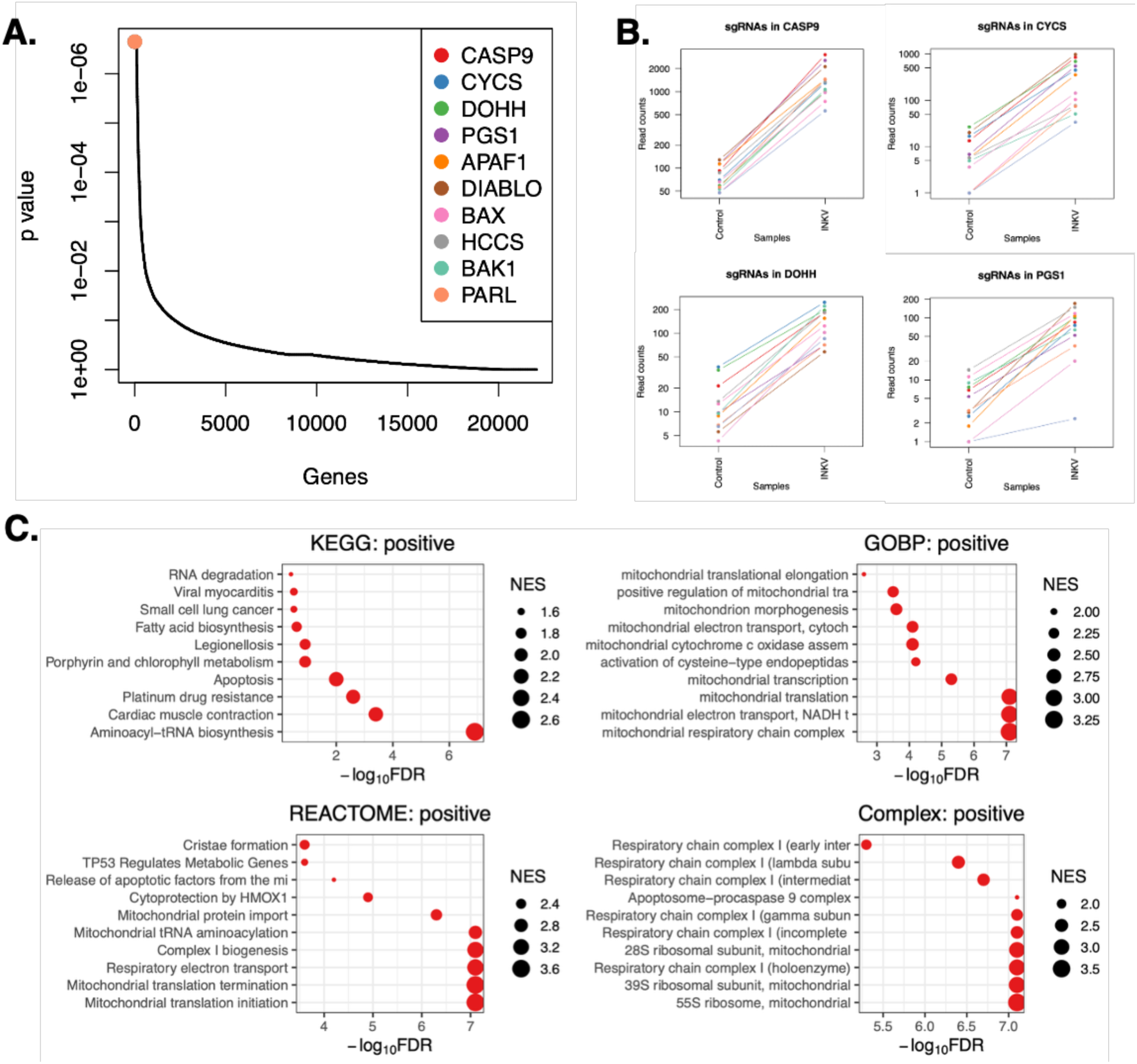
INKV Survival Screen Genetic and Pathway Hits. (A) MAGeCK-VISPR analysis of NGS results from the INKV whole genome knockout survival screen were plotted as a distribution of *p* values across genes, with the top ten hits highlighted. (B) Read count plots from the top four ranked hits from the INKV Screen show nearly universal enrichment from control to INKV samples; each line represents one sgRNA from one replicate. (C) Pathway analysis of genetic hits was conducted using the KEGG, GOBP, Reactome, and Complex databases through the MAGeCK-VISPR platform. Negative log_10_ FDR were plotted for pathway hits with point size corresponding to relative NES.

### No single membrane protein receptor identified

There are currently no known obligate entry receptors for any of the orthobunyaviruses. For both the LACV and INKV whole genome knockout screens, single gene knockouts were insufficient to confer complete immunity to virus-induced cell death. Furthermore, none of the screen top ranked hits were membrane protein receptors that could be candidates for exclusively mediating viral entry (Supplemental Tables 3&4). To further probe for potential receptor proteins, we conducted a targeted CRSIRP membrane protein knockdown screen in BE(2)-C cells with LACV MOI = 1 as the selection pressure. Quality control measures confirmed function of the constitutively expressed dCas9-KRAB enzyme in the BE(2)-C library cells as well as balanced representation of membrane library sgRNAs (data not shown). At the time of genomic DNA harvest, 72 hours post infection, there was 28.2% of survival infected cells. MAGeCK pipeline analysis of the surviving cells revealed no single genes that hit the threshold of FDR < 0.05 to quality as a significant hit. Although some *p* values were below 0.05, these genes did not have compelling enrichment of their sgRNAs across replicates. Ultimately, the membrane protein knockdown LACV survival screen in neuroblastoma cells had null results (Supplemental Figure 4, Supplemental Table 5). This suggests that LACV does not rely on the expression of a single membrane protein receptor to enter cells.

### Small molecule impairment of apoptosis and oxidative phosphorylation prolonged infected cell survival

To confirm the screen results, we aimed to replicate the prolonged survival phenotype of impaired metabolic and apoptotic pathways by using orthogonal manipulations in non-genetically modified cells. To evaluate the effects of disrupting metabolism, we treated cells with oligomycin. Oligomycin inhibits the action of ATP synthase thus thwarting oxidative phosphorylation leading to impaired metabolism [61]. We evaluated cells at 48- and 96-hour timepoints to allow for sufficient survival of untreated wild type cells for comparison, based on our results from the survival screens. Treatment of BE(2)-C wild type cells with 5 μM oligomycin resulted in improved survival for LACV MOI = 1 at 48 hours post infection and for INKV MOI = 1 at 96 HPI (Figure 4A-B). Treatment with oligomycin did not cause phenotypic changes in the cells (Supplemental Figure 1F). Our results show that impairment of mitochondrial energy production provided a survival advantage for virally infected cells, validating our findings from the survival screen.

**Figure 4.**
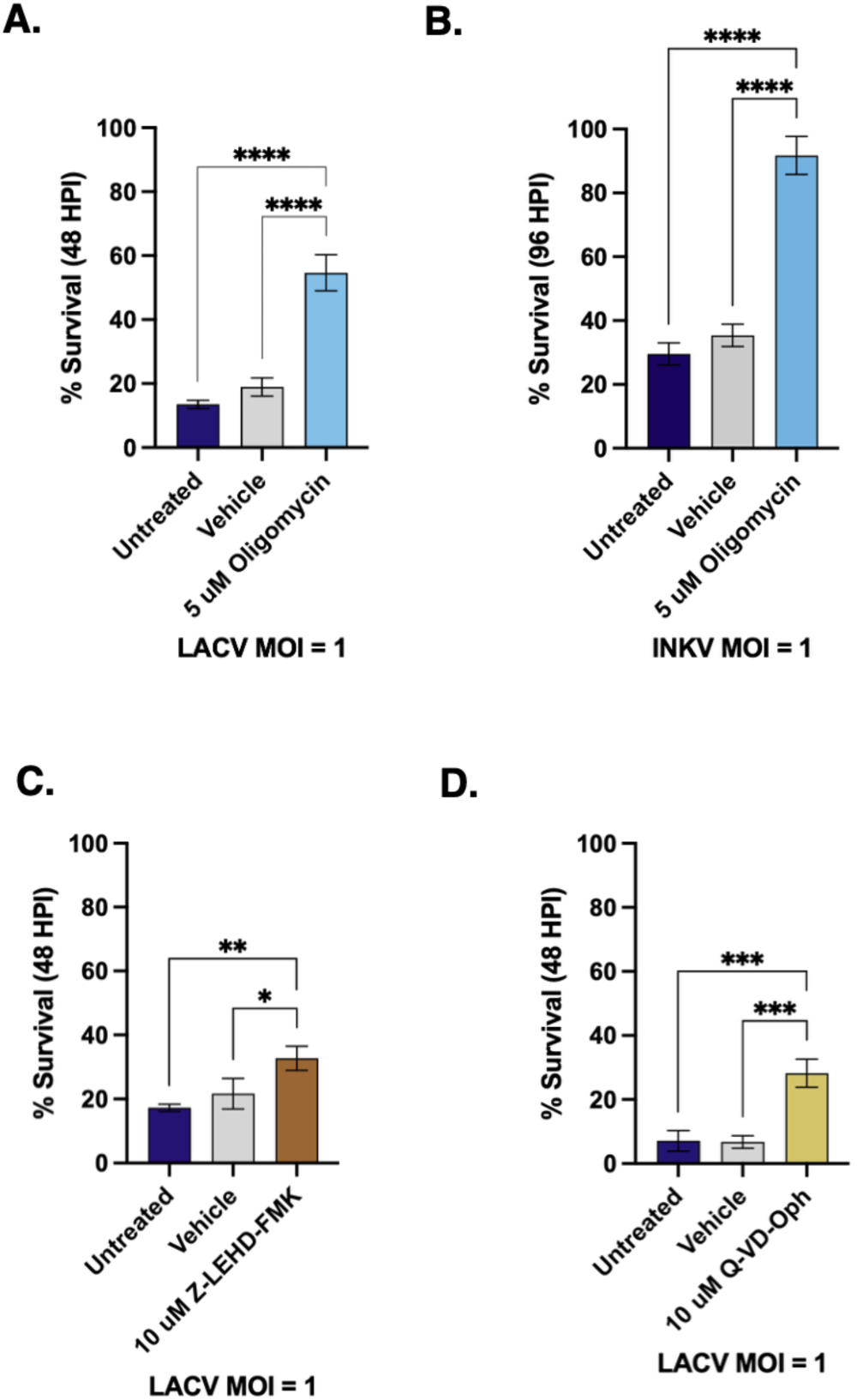
Validation of Metabolic and Apoptotic Screen Hits. (A-B) BE(2)-C wild type cells were treated with nothing, DMSO vehicle, or 5 µM oligomycin ATP synthase inhibitor immediately prior to inoculation with LACV (MOI = 1) or INKV (MOI = 1). At 48 (LACV) or 96 (INKV) HPI, the percent survival was determined as compared to uninfected controls. (C) BE(2)-C WT cells were treated with nothing, DMSO vehicle, or 10 µM Z-LEHD-FMK caspase 9 inhibitor immediately prior to inoculation with LACV (MOI = 1). At 48 (LACV) HPI, the percent survival was determined as compared to uninfected controls. (D) BE(2)-C WT cells were treated with nothing, DMSO vehicle, or 10 µM Q-VD-Oph pan caspase inhibitor immediately prior to inoculation with LACV (MOI = 1). At 48 (LACV) HPI, the percent survival was determined as compared to uninfected controls. (A-D) Asterisks represent significance values from one-way ANOVA as follows: *p ≤0.05, **p ≤ 0.01, ***p ≤ 0.001, ****p ≤ 0.0001; n = 3.

Caspase 9, the initiator caspase for intrinsic apoptosis, was the highest ranked hit for both the LACV and INKV survival screens. To evaluate the effects of blocking caspase 9 in non-genetically modified cells, we treated cells with the caspase 9 inhibitor Z-LEHD-FMK. When BE(2)-C wild type cells were treated with 10 μM Z-LEHD-FMK, they had greater survival of LACV MOI = 1 infection at 48 hours compared to untreated and vehicle treated cells (Figure 4C). We were curious about whether this effect was limited to the intrinsic apoptosis initiator caspase, so we tried a pan caspase inhibitor that acts against the initiator and executioner caspases for both intrinsic and extrinsic apoptosis. The pan caspase inhibitor Q-VD-Oph (10 μM) likewise significantly increased survival of LACV MOI = 1 infection at 48 hours in BE(2)-C wild type cells (Figure 4D) [15]. Treatment with pan caspase inhibitor did not cause phenotypic changes in the cells (Supplemental Figure 1E). Our results show that both specific impairment of caspase 9 and a more global impairment of apoptosis were beneficial, indicating apoptosis is an important mechanism of LACV-induced neuronal death.

### Mild hypothermia and feeding galactose both prolonged infected cell survival

With both LACV and INKV survival screens identifying genes critical for energy production in the mitochondria, we wondered if a more global impairment of energy flux through the cell would result in a similar prolonged survival phenotype. It is established that reductions in temperature lead to reduced metabolic flux [24, 51, 118]. Therefore, we tested whether a reduction in incubation temperature from the standard 37°C to 33°C provided a survival benefit. Of note, the incubation temperature was changed at the time of infection, not prior. For LACV MOI = 0.0001, 0.01, and 1 and for INKV MOI = 0.01, 1, and 100, incubation at 33°C provided a significant survival advantage at 48 and 96 HPI, respectively in BE(2)-C wild type cells (Figure 5A-B). Treatment with mild hypothermia did not cause phenotypic changes in the cells (Supplemental Figure 1G). To determine if this reduced temperature protective effect was limited to the California serogroup of Orthobunyaviruses, we also tested temperature effects on survival rates of Bunyamwera virus (BUNV). BUNV, a neuroinvasive orthobunyavirus in the Bunyamwera serogroup, was chosen as an outgroup comparison to determine the generalizability of the interventions. At BUNV MOI = 0.01 and 1, and we found a significant survival benefit at 72 HPI (Figure 5C). Without genetic perturbations or small molecule inhibitors, we were able to modulate total energy flux in the cell by reducing the temperature to promote a greater survival interval for infection by LACV, INKV, and BUNV.

**Figure 5.**
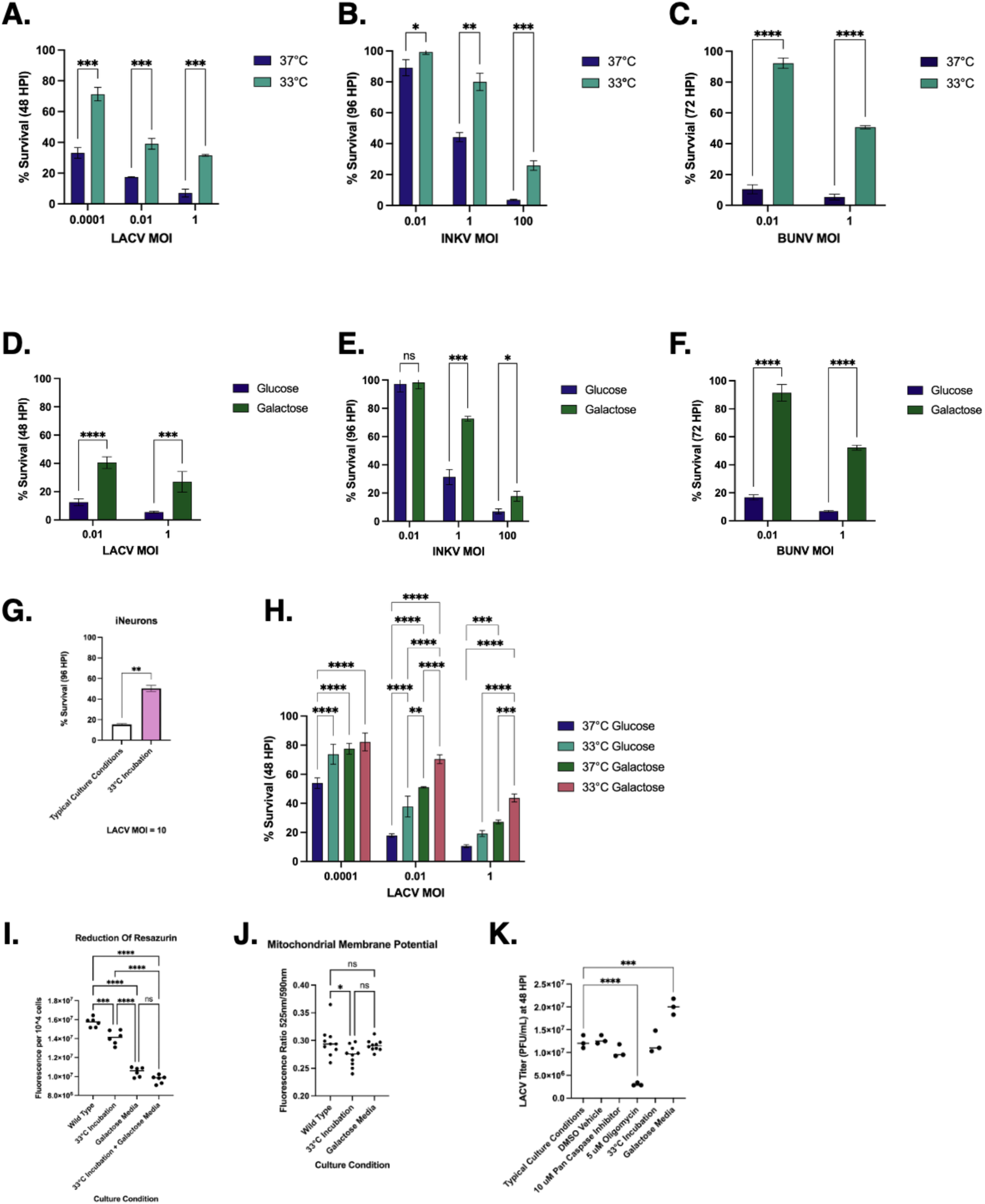
Effect of Metabolic Modulations on Viral Susceptibility. (A-C) BE(2)-C wild type cells were inoculated with LACV (MOI = 0.0001, 0.01, 1), INKV (MOI = 0.01, 1, 100), or BUNV (MOI = 0.01, 1) then incubated at either 37°C or at 33°C. At 48 (LACV), 72 (BUNV), or 96 (INKV) hours post infection (HPI), the percent survival was determined as compared to uninfected controls. (D-F) BE(2)-C wild type cells were pre-grown for at least three days and subsequently maintained in media with either 10 mM galactose or 10 mM glucose as the primary carbon source prior to inoculation with LACV (MOI = 0.01, 1), INKV (MOI = 0.01, 1, 100), or BUNV (MOI = 0.01, 1). At 48 (LACV), 72 (BUNV), or 96 (INKV) HPI, the percent survival was determined as compared to uninfected controls. (G) WTC11-NGN2 iNeurons were cultured until mature at 14 days post-differentiation induction at low confluence on 10-cm plates. They were inoculated with LACV (MOI = 10) and then incubated either 37°C or at 33°C. At 96 hours post infection (HPI), the percent survival was determined as compared to uninfected controls. (H) BE(2)-C wild type cells were pre-grown for at least three days and subsequently maintained in media with either 10 mM galactose or 10 mM glucose as the primary carbon source prior to inoculation with LACV (MOI = 0.0001, 0.01, 1). Cells were then incubated at either 37°C or at 33°C for 48 hours post infection (HPI). Percent survival was determined as compared to uninfected controls. (I) The reduction potential of BE(2)-C wild type cells subjected to baseline culture conditions, 33°C incubation, 10 mM galactose media, or the combination of 33°C incubation plus 10 mM galactose media for two days was measured by the reduction of resazurin live cell dye to resorufin over the course of 2.5 hours. Fluorescence was measured at 560/590 nm and scaled to cell count. (J) The mitochondrial membrane potential of BE(2)-C wild type cells subjected to baseline culture conditions, 33°C incubation or 10 mM galactose media for two days was measured by the ratio of monomeric cytoplasmic (490/525 nm) to aggregated mitochondrial (540/590 nm) JC-10 where a lower ratio indicates higher mitochondrial membrane potential. (K) The viral titer in PFU/mL of the cell culture supernatant was measured at 48 hours post infection of BE(2)-C wild type cells subjected to various treatment conditions and infected with LACV MOI = 1. Titers were performed via vero cell-based plaque assays. (A-K) Asterisks represent significance values from unpaired t-tests (A-G) or ANOVAs (H-K) as follows: *p ≤0.05, **p ≤ 0.01, ***p ≤ 0.001, ****p ≤ 0.0001; (A-H; K) n = 3; (I) n = 6; (J) n = 10.

To assess whether the benefits of impaired metabolism we observed in the BE(2)-C neuroblastoma cells applied to a more clinically relevant neuronal model system, we tested the effects of mild hypothermia on human iPSC derived neurons (iNeurons). For LACV MOI = 10, incubation at 33°C greatly improved survival at 96 hours post infection as compared to incubation at 37°C (Figure 5G). The replication of the survival benefit found in the neuroblastoma cells in the iNeurons is promising for potential clinical translation.

To further manipulate metabolism, we substituted galactose for glucose as the primary carbon source. Per sugar molecule, metabolism of galactose produces less energy and increases reliance on oxidative propargylation as compared to glucose [3, 55, 67, 89, 93, 94, 98]. To allow for the cellular stores of glucose to be processed and depleted prior to infection, cells for these experiments were pre-cultured in 10 mM galactose media for at least 3 days prior to viral inoculation. For LACV MOI = 0.01 and 1, and for INKV MOI = 0.01, 1, and 100, swapping 10 mM glucose for 10 mM galactose as the primary carbon source provided a significant survival benefit at 48 and 96 HPI, respectively in BE(2)-C wild type cells (Figure 5D-E). Treatment with galactose media did not cause phenotypic changes in the cells (Supplemental Figure 1H). Once again, we tested whether this benefit would extend to BUNV; we found that for BUNV MOI = 0.01 and 1 at 72 HPI there was a significant survival benefit of galactose media (Figure 5F). Thus, reducing the total energy per molecule of carbon catabolized increased the survival interval for viral infection.

Our results showed that both reducing temperature and subbing galactose for glucose resulted in increased cell survival, however neither method resulted in full protection of cells. We therefore were interested in determining whether the 33°C incubation temperature and the galactose carbon source interventions were operating by the same, or distinct, mechanisms. To see if there was an additive benefit, we tested the effect of combinations of 33°C incubation and galactose media on the survival proportion of BE(2)-C wild type cells infected with LACV MOI = 0.0001, 0.01, and 1 at 48 HPI. We found that the combination of 33°C incubation and galactose media provided a stronger survival benefit than either intervention applied in isolation (Figure 5H). Thus, the interventions either failed to saturate a shared mechanism or operated by different mechanisms.

Beyond the genetic manipulations of the knockout screens and the small molecule inhibitors targeting ATP synthase or apoptotic caspases, culture condition modifications to reduce total energy in the cell provided a prolonged survival benefit for LACV and INKV. Interestingly, both mild hypothermia and galactose substitution, as well as the combination of the two, slowed the division rate of BE(2)-C wild type cells substantially: likely another downstream effect of the energy impairment that offers viral protection (Supplemental Figure 5). Together, these results suggest that the host cell energetic flux is critical in modulating susceptibility to virus-induced cell death.

### Host cell energetics regulate viral susceptibility

Since we expected our interventions would result in lower total energy availability in the cell, we tested the total reducing power of wild type, non-infected cells grown in typical culture conditions, 33°C incubation, galactose media, or the combination of 33°C incubation and galactose media. The reducing power of a cell is a measure of the energy readily available to complete cellular processes or to an invading virus. A higher reducing power indicates a more biosynthetic or anabolic state. The reduction of resazurin, a non-toxic cell-permeable dye is a convenient measure of total cellular reducing potential. As measured by the reduction of resazurin to resorufin, typical culture conditions resulted in the highest reducing power, 33°C incubation had an intermediate reducing power, and both galactose media and the combination of 33°C incubation plus galactose media had the lowest reducing power (Figure 5I). While both interventions on their own lowered reducing power, the combination of the two did not perform significantly better than galactose media in isolation. The additive survival benefit of combining 33°C incubation with galactose media cannot be attributed to a lower cellular reducing power (Figure 5I). Overall, the loss of reducing power in any of these culture conditions could limit the energy available for viral processes in the cell, thus lowering the cytotoxicity and prolonging survival.

Given that impairment of mitochondrial energy production was the strongest pathway hit from both screens, we tested how 33°C incubation and galactose media altered the mitochondrial membrane potential. JC-10 is a cationic, fluorescent dye that forms longer wavelength emitting aggregates in polarized mitochondria. Based on the wavelength emission of JC-10, we found that 33°C incubation had a greater mitochondrial membrane potential than typical culture conditions, but galactose media did not significantly differ from either 33°C incubation or typical culture conditions (Figure 5J). This indicates that if 33°C incubation or galactose media are operating through the mitochondria, they are doing it in such a way that leaves the mitochondrial membrane potential intact. In sum, the reducing power but not necessarily the mitochondrial membrane potential of the host cell affect the vulnerability of that cell to virus-induced cell death.

### Mild hypothermia and feeding galactose do not reduce viral titer

To assess the effect of each of the interventions on the production of infectious viral particles released into the cell culture media, we performed a plaque assay titer of culture supernatants at 48 hours post infection of wild type BE(2)-C cells infected with LACV MOI = 1. Supernatants were titered on Vero cells. The viral titers for DMSO vehicle, 10 µM pan caspase inhibitor, and 33°C incubation were not significantly different than no intervention or typical culture conditions. Interestingly, galactose media increased the viral titer significantly, and 5 µM oligomycin treatment reduced the titer significantly (Figure 5K). This suggests that for 33°C incubation and 10 µM Pan Caspase inhibitor, the survival benefit is not due to a reduction in the production of infectious viral particles. At least part of the 5 µM oligomycin survival benefit may be due to reduction in virus production. For galactose carbon source substitution, the survival benefit occurs despite an increased production of virions. These results suggest that there are a variety of mechanisms to protect neuronal cells from death from LACV, and they can be independent from reducing virus production.

### Feeding galactose but not mild hypothermia increased survival via reduction in caspase 9 dependent apoptosis

To assess the degree to which 33°C incubation and galactose media may be acting through an impairment of apoptosis, we first tested them in combination with the pan caspase inhibitor Q-VD-Oph. Treatment with 10 μM Q-VD-Oph was shown to largely obliterate caspase 9 activation when challenged with either LACV MOI = 1 or the apoptosis-inducing camptothecin at 4 ug/mL (Figure 6A). Thus, if there is a further benefit of any intervention when employed in combination with 10 μM Q-VD-Oph, it is likely not operating through impairment of caspase 9 dependent apoptosis.

**Figure 6.**
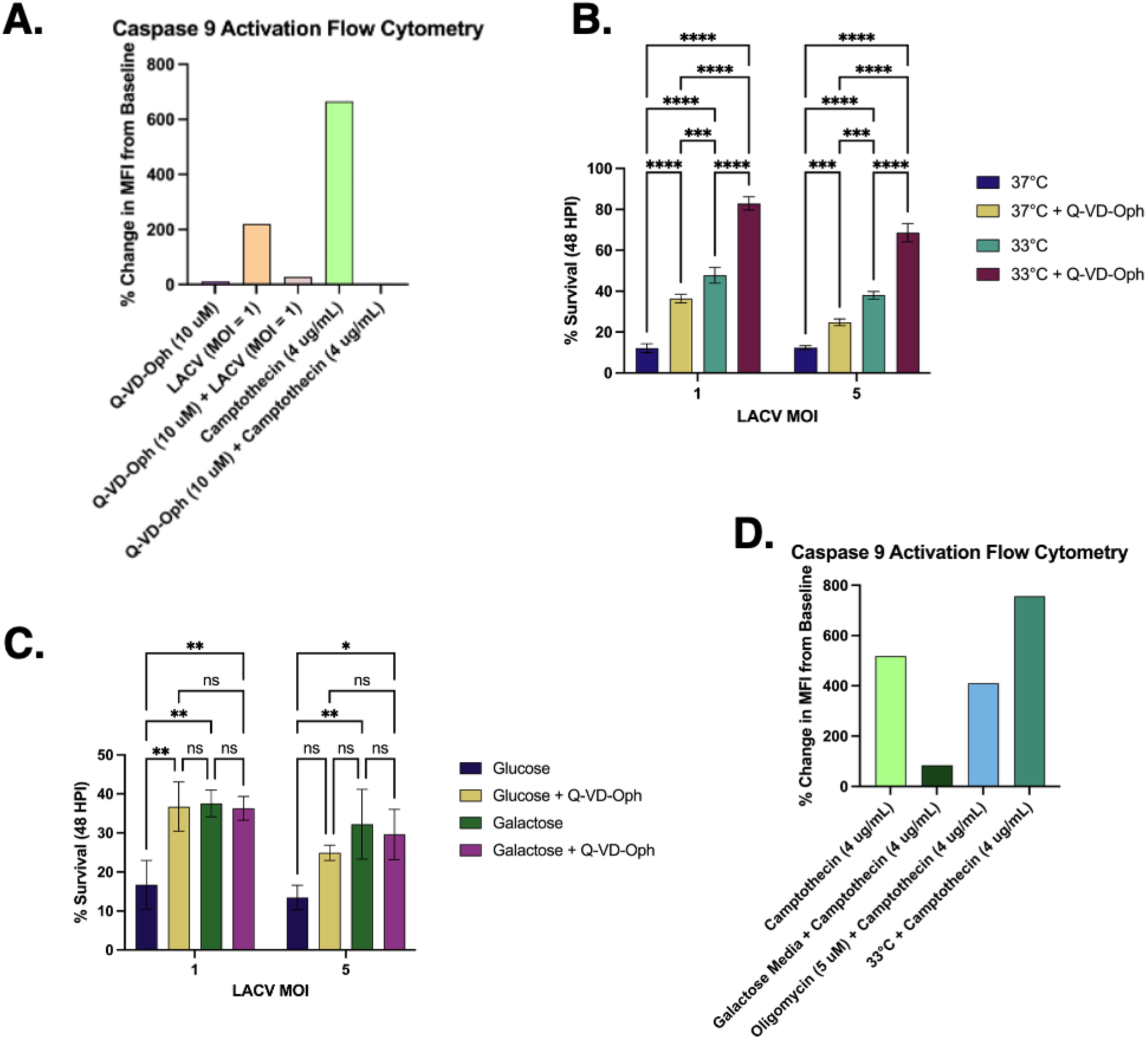
Effect on Metabolic Modulations on Apoptotic Suppression. (A&D) The effect of interventions on apoptosis was interrogated by measuring activated caspase 9 approximately 24 hours post treatment via flow cytometry. Camptothecin (4 ug/mL) was utilized to induce apoptosis via caspase 9 action. Activated caspase 9 was detected using a fluorochrome inhibiter of caspases (FLICA) conjugated to a green FAM fluorescent probe. Samples were run on a CytoFLEX LX cytometer and the percent change in median fluorescence intensity (MFI) from BE(2)-C wild type, untreated cells was reported. (A) 10 µM Q-VD-Oph pan caspase inhibitor was challenged with LACV MOI = 1 or camptothecin (4 ug/mL). Each bar represents the MFI from a single run of flow cytometry; traces are available in Supplemental Figure 6. (B) BE(2)-C wild type cells were treated with DMSO vehicle or 10 µM Q-VD-Oph pan caspase inhibitor immediately prior to inoculation with LACV (MOI = 1, 5) then incubated at either 37°C or at 33°C for 48 HPI. Percent survival was determined as compared to uninfected controls. (C) BE(2)-C wild type cells were pre-grown for at least three days and subsequently maintained in media with either 10 mM galactose or 10 mM glucose as the primary carbon source. Cells were treated with either DMSO vehicle or 10 µM Q-VD-Oph pan caspase inhibitor immediately prior to inoculation with LACV (MOI = 1, 5). At 48 HPI, percent survival was determined as compared to uninfected controls. (B-C) Asterisks represent significance values from unpaired t-tests as follows: ns = not significant, *p ≤0.05, **p ≤ 0.01, ***p ≤ 0.001, ****p ≤ 0.0001; n=3. (D) 10 mM galactose as the primary carbon source (initiated at least 3 days prior to and maintained through inoculation and incubation), 5 µM oligomycin, and 33°C incubation temperature were challenged with camptothecin (4 ug/mL). Each bar represents the MFI from a single run of flow cytometry; traces are available in Supplemental Figure 6.

When 33°C incubation was tested in combination with 10 μM Q-VD-Oph for LACV MOI = 1 and 5 (a higher MOI was included to increase the apoptotic induction signal per cell and allow for greater discernment of potential effect), there was an additive benefit such that the combination provided a greater survival benefit for BE(2)-C wild type cells at 48 HPI than either employed in isolation (Figure 6B). The opposite pattern was seen for galactose media tested in combination with 10 μM Q-VD-Oph for BE(2)-C wild type cells infected with LACV MOI = 1 and 5 at 48 HPI; there was no additional benefit to the combination (Figure 6C). The data suggests that galactose media is operating through shared anti-apoptotic mechanisms with Q-VD-Oph while 33°C is not. This could account for the survival benefit of the galactose media despite the increase in viral titer compared to glucose media (Figure 5K).

To follow-up on this finding, we employed flow cytometry assay that detects active caspase 9. Uninfected BE(2)-C wild type cells were treated with 4 ug/mL of camptothecin to induce caspase 9 activity. Galactose media, 5 μM oligomycin, and 33°C incubation were tested against the action of camptothecin to see if they could mitigate or obliterate the induced caspase 9 activity. As previously shown, 10 μM Q-VD-Oph did obliterate the action of 4 ug/mL camptothecin (Figure 6A). Galactose media provided the sharpest reduction in caspase 9 activity, supporting the earlier finding that it likely acts through impairment of apoptosis (Figure 6D). Treatment with 5 μM oligomycin and 33°C incubation had no such mitigating effect on caspase 9 activation (Figure 6D). Flow traces, median fluorescence intensities, median absolute deviations, and cell counts for each treatment are shown in Supplemental Figure 6. In summary, while both galactose media and 33°C incubation lower the reducing potential of a cell and provide a prolonged survival benefit for viral infection, only galactose seems to be acting through impairment of apoptotic processes. Oligomycin treatment seems to be operating through reduction of viral titer (Figure 5K).

## Discussion

To our knowledge, we have reported the first genome-wide CRISPR survival screens for CSG viruses. For both a highly pathogenic virus and a largely benign virus, we found that host cell mitochondrial energy production was a primary positive regulator of cell death. We expanded on this finding by demonstrating survival advantages for infected cells that had their energy production modulated by either mild hypothermia or by feeding galactose as the primary carbon source. Furthermore, we were able to generalize these findings by replicating the survival advantage for infection with BUNV, a more distantly related orthobunyavirus.

Despite their disparate clinical phenotypes, both the single gene hits and enriched pathways were overwhelmingly shared between LACV and INKV (Figure 1). This alignment suggests that host cell-virus interactions likely are not the cause of the varied pathogenicity in humans – other studies have found differences in viral replication rates and evoked immune responses that may be more culpable [4, 8, 9, 26–28, 44, 69, 86, 100, 111, 113–115]. General interventions that target shared host cell pathways, however, have the potential to be useful for orthobunyaviruses generally and for any newly emergent, reassorted viruses.

Interestingly, no single gene knockout or knockdown was sufficient to entirely prevent cell death, only delay it (Supplemental Tables 3-5). There are several primary interpretations of this result: (1) A complete knockout of the specific (and unknown) membrane protein receptor is required to provide a survival benefit or (2) the virus is utilizing multiple redundant entry receptors such that impairment of a single route is not protective against infection or (3) the virus is using entry methods that do not rely on a protein membrane receptor. Given that a single membrane receptor was not a top hit in the knockout whole genome screens, we suspect that LACV is utilizing redundant methods of access. These methods of access may be facilitated by interaction with the low-density lipoprotein receptor-related protein 1 (LRP1) which has recently been shown to augment LACV infection [21, 38, 40, 41].

As expected, pro-apoptotic machinery was the top hit in both screens. Highly ranked pro-apoptotic genes include CASP9, DIABLO, CYCS, APAF1, BAK1, HCCS, and PARL (Supplemental Tables 3&4). This finding was validated with the use of both a specific caspase 9 inhibitor and a pan caspase inhibitor, which increased survival of LACV infection (Figure 4C-D). As this finding aligns with the literature, it serves as a good internal control for screen validity [18, 80, 114]. Our major novel finding was the shared reliance on respiratory energy production: GOBP pathway analysis found enrichment in mitochondrial respiratory chain complexes, mitochondrial electron transport, mitochondrial translation, and cellular respiratory; REACTOME analysis found enrichment in mitochondrial translation termination, mitochondrial translation initiation, complex 1 biogenesis, and respiratory electron transport; and Complex analysis found enrichment in respiratory chain complex 1, 39S ribosomal subunit mitochondrial, and 55S ribosome mitochondrial (Figures 2&3). In essence, cells with impaired metabolism survived CSG infection longer than those with fully functional oxidative phosphorylation, and survival was shown to be both caspase dependent and independent.

The link between impaired mitochondrial function and improved survival was validated by oligomycin treatment, a small molecule inhibitor of ATP synthase (Figure 4A-B). Cells with a reduced energy flux were more resistant to viral cell death, a finding that perhaps plays a role in the clinical propensity for LACV encephalitis to preferentially affect children. LACV-E is both more prevalent and more severe in younger patients [13, 19, 20, 39, 70, 77, 78, 106]. Research in the field of aging has found that older individuals have impaired energy homeostasis in the central nervous system, and the peak age range for brain glucose uptake and metabolism (1-7 years old) correlates with the peak age range for susceptibility to LACV-E [43, 59, 68, 74]. It could be that the increased energy available in a developing brain allows for a more pathogenic viral infection. A previously identified antiviral, rottlerin, that likewise increases survival of infected neuroblastoma cells in culture functions by reducing ATP production in the host cell further bolsters this energy flux link [76].

We were interested in clinically relevant extensions of the observation that impairment of energy production prolonged survival, so we sought to determine if we could increase survival by lowering incubation temperature in culture (therapeutic hypothermia) and feeding cells galactose rather than glucose (carbon source selection). In this study, we tested a reduced temperature of 33°C because this temperature has been used in human patients in therapeutic hypothermia protocols [57, 58, 99, 109, 120]. We found a marked survival benefit for cells incubated at 33°C that held true for LACV, INKV, and BUNV as well as for neuroblastoma cells and human induced neurons (Figure 5). We also tested the effect of replacing glucose in cell culture media with galactose as the primary carbon source. This replacement causes an increase in reliance on oxidative phosphorylation, depriving cells of much of the energy typically received from glycolysis, particularly in cancer cell lines [3, 55, 67, 89, 93, 94, 98]. Notably, this diverges a bit from the screen findings that focused on impaired mitochondrially based energy production, largely altering cytoplasmic energy metabolism instead. Once again, we found a marked survival benefit for cells fed galactose for LACV, INKV, and BUNV (Figure 5), suggesting the benefit may be due to total energy flux reduction rather than specifically mitochondrial energy flux reduction. The generalized benefit regardless of orthobunyavirus identity posits carbon source changes and therapeutic hypothermia as promising potential clinical treatments, even in cases of encephalitis where the viral agent is not immediately diagnosable.

Feeding galactose and mild hypothermia had an additive effect, such that infected neuroblastoma cells cultured in 33°C and fed galactose had the highest survival percentages. One potential mechanism we identified for both mild hypothermia and feeding galactose is a decrease in total energy available in the cell, as measured by reducing potential (Figure 5). Neither decreased virion production, with galactose media paradoxically increasing it. The increase in viral titer for galactose media was surprising, but we hypothesize it could be due to an increased production or availability of metabolites useful for virion production. Increasing the reliance on oxidative phosphorylation likely shifted the metabolism homeostasis of the cell, potentially allowing for more free lipids, nucleic acids, or amino acids for virion building, but this remains speculative.

To determine the extent to which feeding galactose and mild hypothermia could be causing a reduction in apoptosis, we tested them each in the presence of pan caspase inhibitor (titrated to eliminate caspase 9 induction from LACV infection). Interestingly, mild hypothermia had an additive benefit to the pan caspase inhibitor while feeding galactose had no further benefit compared to the pan caspase inhibitor (Figure 6). A follow-up study using camptothecin as an apoptosis inducer again showed that galactose media acted by reducing caspase 9-dependent apoptosis while mild hypothermia did not (Figure 6). Thus, feeding galactose could be decreasing the available ATP necessary to power apoptotic induction as demonstrated by impairment of caspase 9-dependent apoptosis, which would account for improved cell survival despite increased virion production, whereas mild hypothermia appears to be reducing neurotoxicity by a different, non-apoptotic mechanism. We did not determine this mechanism experimentally, but possibilities include: impairment of viral entry to the cell, down regulation of transcription and/or translation, alteration in viral and host protein conformations, changes in membrane fluidity, modified timing of cell cycle progression, global decrease in endocytic and exocytic trafficking, variability in infectable cell status, and alteration of catabolic processes, among others [10, 12, 17, 23, 29, 30, 34, 50, 62, 82, 85, 87, 105, 107, 108, 117].

Substituting galactose as the primary carbon source and incubating at a mildly hypothermic temperature had striking survival benefits not only for neuroblastoma cells infected with the clinically disparate LACV and INKV, but also for cells infected with the more distantly related orthobunyavirus BUNV. Furthermore, mild hypothermia had a protective effect in human induced neurons. Therapeutic hypothermia, which consists of cooling the core body temperature of human patients to about 32-34°C, has been used clinically for the treatment of ischemic brain injuries [2, 5, 64, 79, 119]. Recent small-scale studies have investigated its use for space-occupying lesions or otherwise increased intracranial pressures in the setting of encephalitis [57, 58, 99, 109, 120]. In this study, we employed mild hypothermia in cell culture, so the benefit could not be attributed to mitigating space-occupying lesions (which cause increased pressure in the brain) and was isolated to metabolic effects. It is possible, therefore, that the benefit could be additive in patients where therapeutic hypothermia treats both space-occupying lesions and reduces cellular metabolism.

Intravenous galactose therapy is typically used clinically to aid in the diagnosis of liver impairment and inborn errors of metabolism [56, 72, 95, 101, 102, 104]. In this study, we used galactose-based cell culture media to reduce total energy flux and impair caspase 9-dependent apoptosis. Further investigations should test whether IV galactose therapy could provide similar benefits for patients infected with arboviruses. Both substituting galactose as the primary carbon source and therapeutic hypothermia hold promise for future, non-invasive clinical treatments for these, and perhaps, other viral encephalitides.

In sum, we have reported two whole genome knock-out survival CRISPR screens for LACV and INKV. We identified metabolic impairment as a protective factor for host cell survival and demonstrated several interventions to manipulate host cell metabolism to increase survival. With LACV-E a current public health threat and the emergence of more virulent, widespread CSG viruses a future public health threat, this study provides an opportunity to better understand how these viruses interact with host cell machinery to cause neuronal death. It also posits therapeutic hypothermia and carbon source (dietary) interventions as plausible treatments.

