## Supplementary Figures for "Energy Flux Regulates Cell Death Induced by California Serogroup Orthobunyaviruses"

**Supplemental Figures**

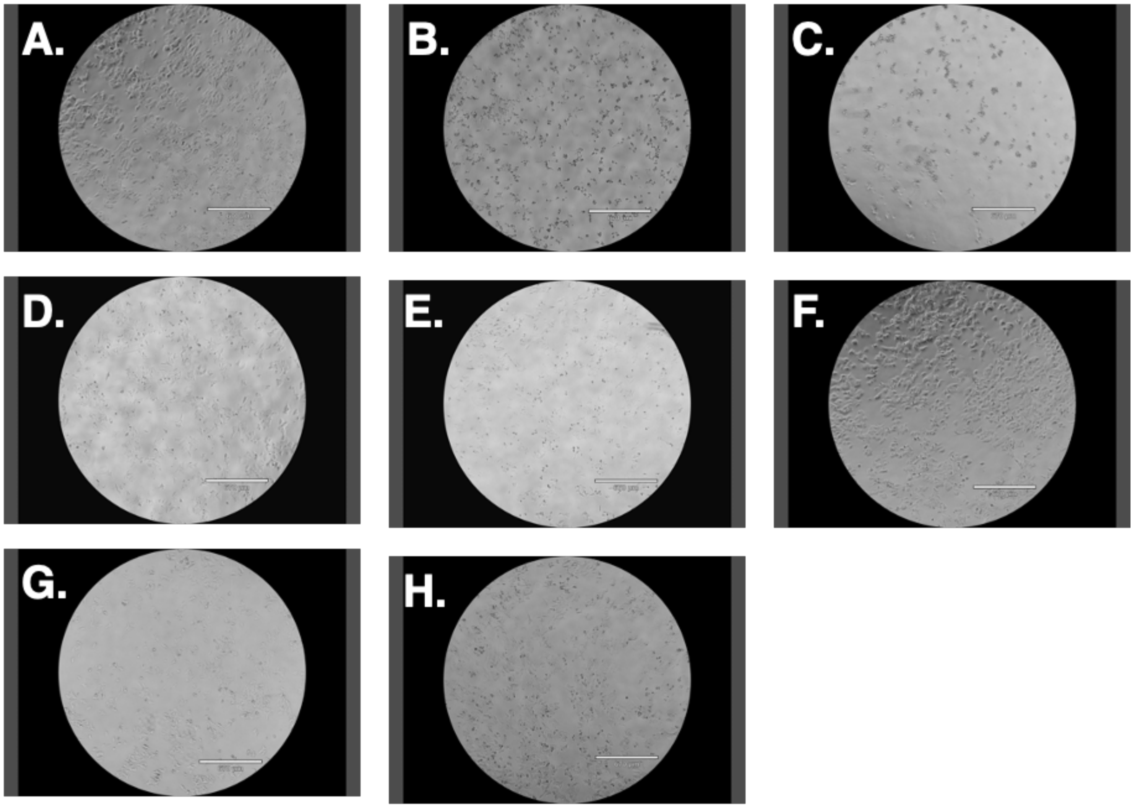


Supplemental Figure 1. Neuroblastoma cell culture with LACV cytotoxicity and treatment conditions.

(A) BE(2)-C neuroblastoma wild type cells under typical culture conditions showed typical morphology then changed to show progress cytopathic effect (CPE) at 24 hours post infection (B) and 48 hours post infection (C) with LACV MOI = 1. None of the treatment conditions caused a change in the typical BE(2)-C neuroblastoma wild type morphology as shown: (D) DMSO vehicle, (E) pan caspase inhibitor, (F) oligomycin, (G) 33°C incubation, and (H) galactose media.

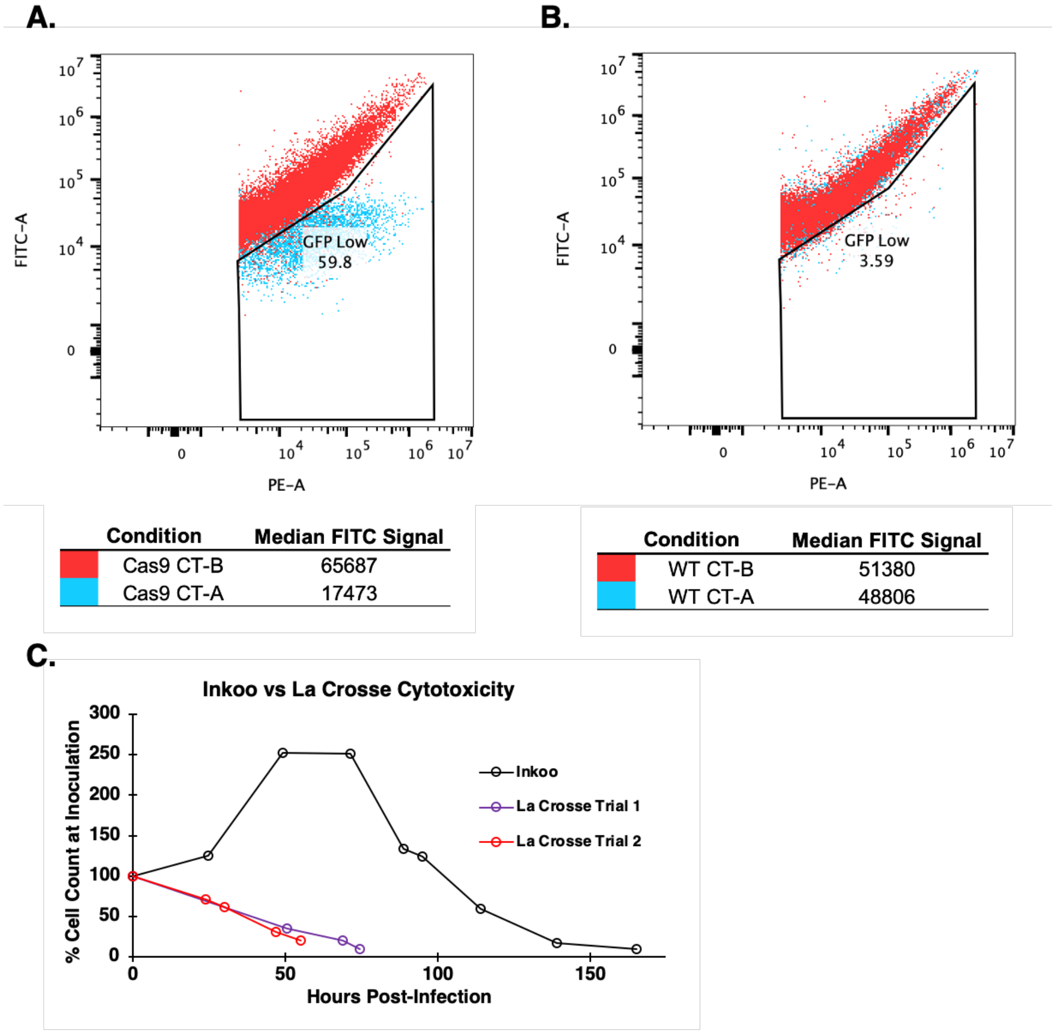


Supplemental Figure 2. Preparation for Survival Screens

(A-B) The function of the Cas9 enzyme was verified prior to screening using sgRNA targeting fluorescent probes from a commercial kit (Cellecta, catalog #CRUTEST). Briefly, BE(2)-C monoclonal Cas9 expressing cells (A) and BE(2)-C wild type cells (B) were transfected with either CT-A, a lentivirus vector containing red fluorescent protein (RFP), green fluorescent protein (GFP), and a sgRNA targeting GFP OR CT-B, a lentivirus vector containing RFP, GFP, and a non-targeting sgRNA. Function of Cas9 was assayed as the decrease in GFP signal for CT-A transfected, RFP-positive cells compared to CT-B transfected, RFP-positive cells. (C) The duration of viral infection required to achieve about a 10% survival of BE(2)-C Cas9 Brunello library cells was determined by inoculation with LACV or INKV (both MOI = 1) and serial cell counts.

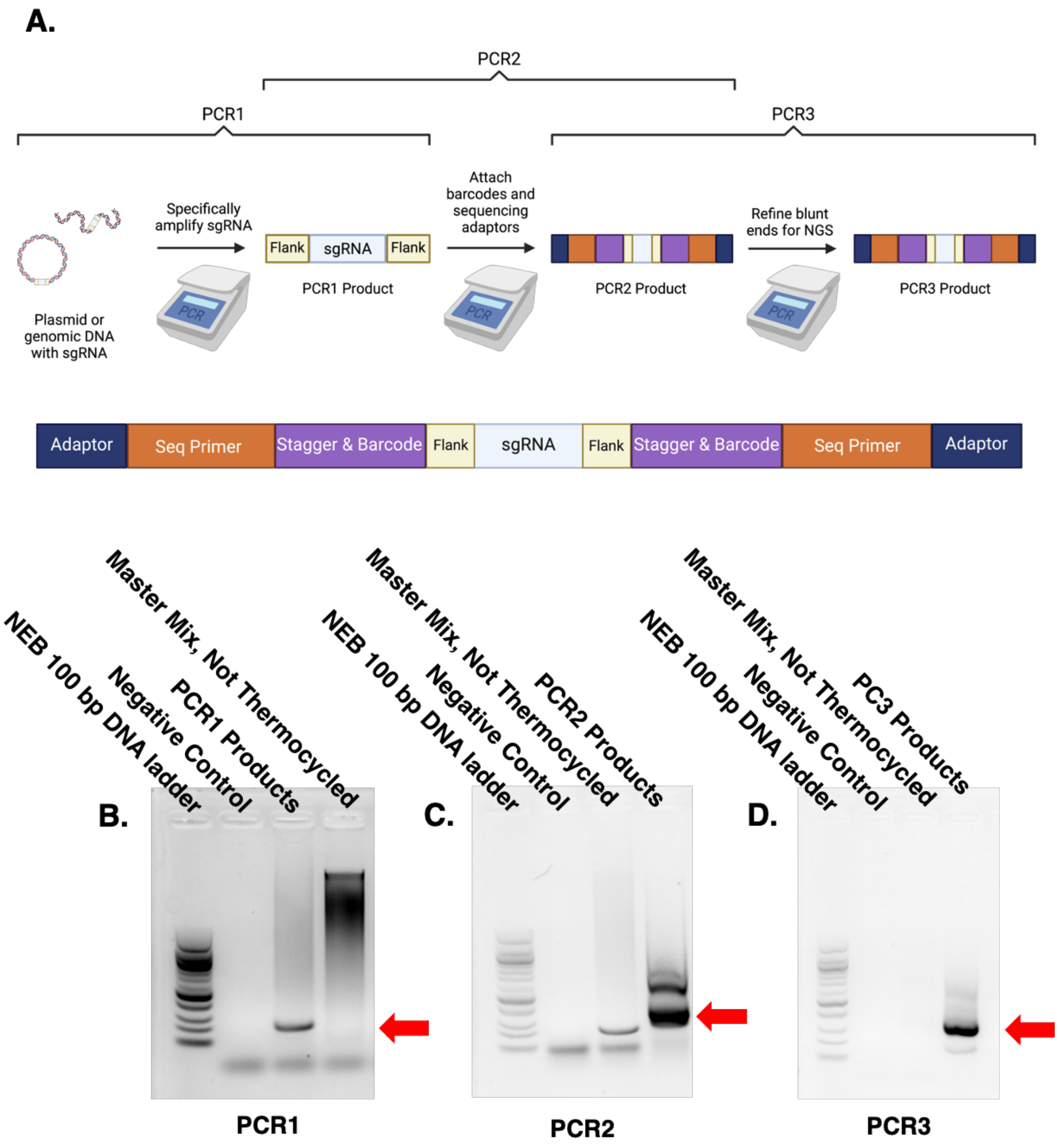


Supplemental Figure 3. PCR Preparation from Genomic DNA to NGS

(A) PCR1 specifically amplified the sgRNA and flanking regions from either plasmid or genomic DNA. PCR2 attached barcodes, stagger sequences, and Illumina sequencing adaptors, and PCR3 refined the blunt ends to improve NGS outcomes. All primers are detailed in Supplemental Table 2. (B-D) Representative DNA gels showing PCR products for PCR1 (B), PCR2 (C), and PCR3 (D). The PCR3 product was subsequently subjected to gel extraction and ready for NGS. (B-D) Red arrow indicates desired product.

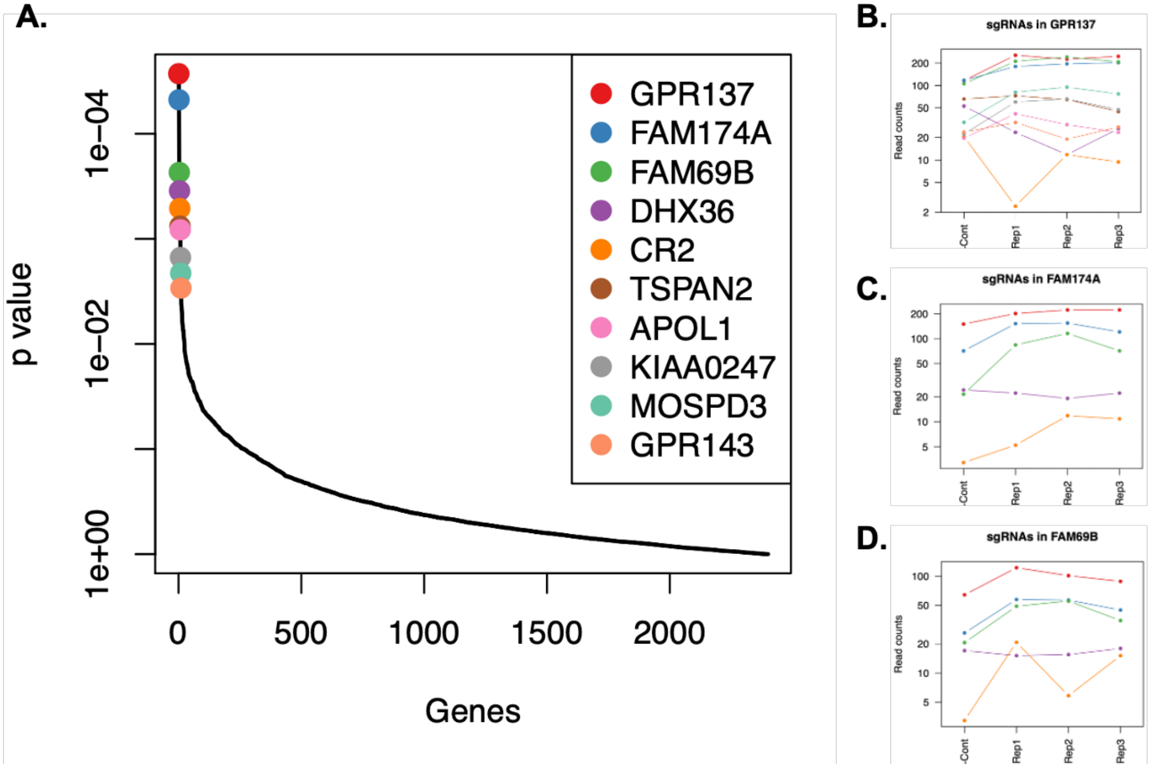


Supplemental Figure 4. LACV Membrane Protein Receptor Knockdown Survival Screen

(A) MAGeCK-VISPR analysis of NGS results from the LACV targeted membrane protein knock down survival screen were plotted as a distribution of *p* values across genes, with the top ten hits highlighted. (B-D) Read count plots from the top four ranked hits from the LACV Screen show very minimal change in sgRNA read counts between the library control and the virally selected cells.

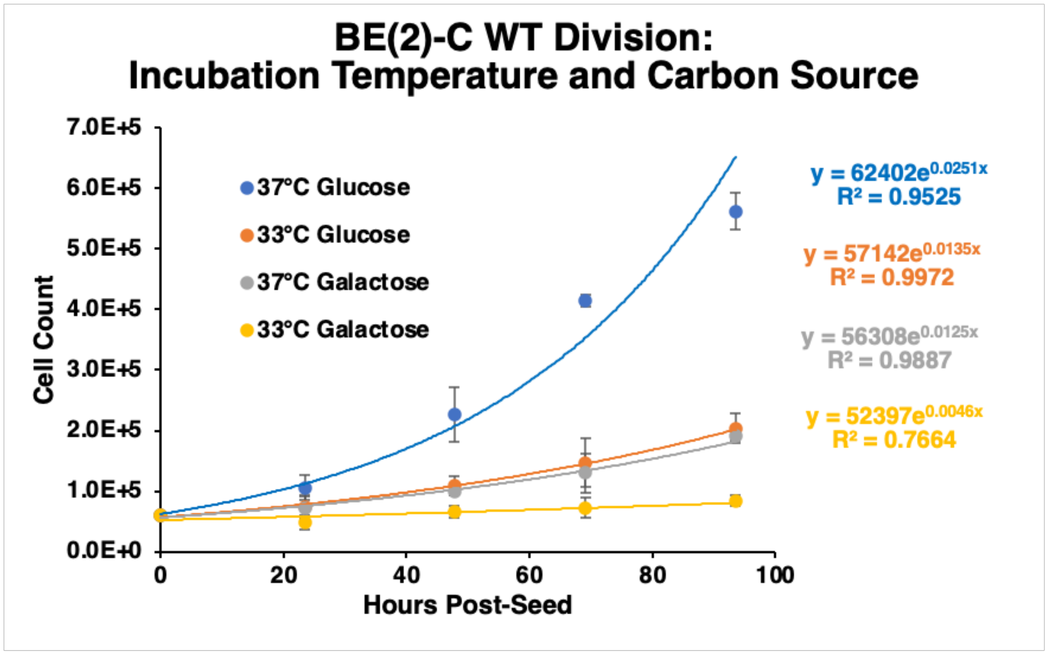


| Cell Culture Condition | Doubling Time (Hours) |
|---|---|
| 37°C Glucose | 27.6 |
| 33°C Glucose | 51.3 |
| 37°C Galactose | 55.5 |
| 33°C Galactose | 150.7 |

Supplemental Figure 5. Effect of Temperature and Carbon Source on Cell Division Rate

BE(2)-C wild type cells were cultured either in media containing 10 mM galactose or 10 mM glucose as the primary carbon source and incubated at either 33ºC or 37ºC with serial cell counts; n=3. Doubling times (in hours) were calculated based on exponential lines of fit.

**A.**

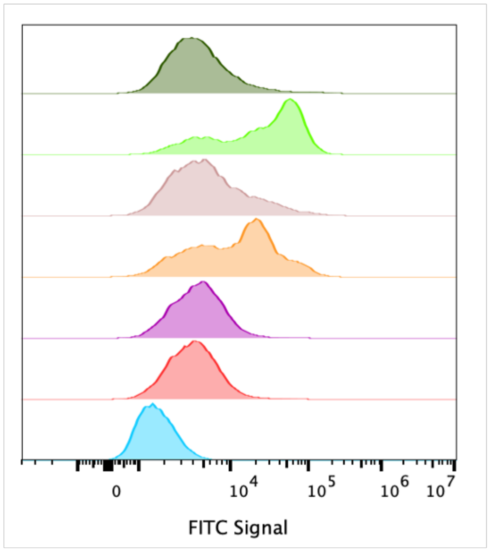


| Condition | Median Fluorescence Intensity | Median Absolute Deviation | Cell Count |
|---|---|---|---|
| Unstained, no treatment | 1,594 | 520 | 22,827 |
| No treatment | 3,905 | 1,432 | 26,436 |
| Q-VD-Oph (10 uM) | 4,405 | 1,661 | 22,183 |
| LACV (MOI = 1) | 12,498 | 8,619 | 18,998 |
| Q-VD-Oph (10 uM) + LACV (MOI = 1) | 5,016 | 2,517 | 19,333 |
| Camptothecin (4 ug/mL) | 29,860 | 22,920 | 21,545 |
| Q-VD-Oph (10 uM) + Camptothecin (4 ug/mL) | 4,111 | 1,652 | 19,746 |

**B.**

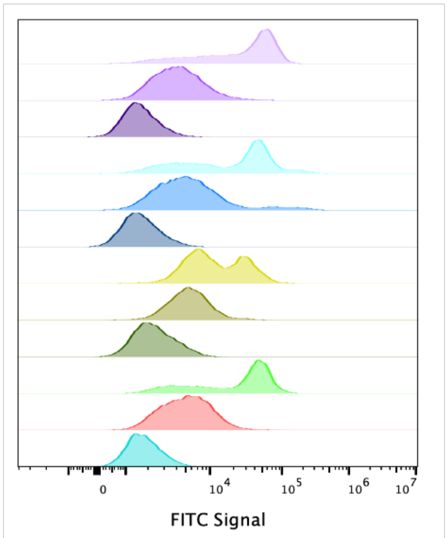


| Condition | Median Fluorescence Intensity | Median Absolute Deviation | Cell Count |
|---|---|---|---|
| Unstained, No Treatment | 1,716 | 528 | 14,051 |
| No Treatment | 5,250 | 2,306 | 21,165 |
| Camptothecin (4 ug/mL) | 32,395 | 22,353 | 22,783 |
| Unstained, Galactose Media | 2,229 | 760 | 13,457 |
| Galactose Media | 5,567 | 1,891 | 14,154 |
| Galactose Media + Camptothecin (4 ug/mL) | 10,269 | 5,566 | 17,418 |
| Unstained, Oligomycin (5 uM) | 1,576 | 541 | 20,086 |
| Oligomycin (5 uM) | 5,184 | 2,506 | 20,108 |
| Oligomycin (5 uM) + Camptothecin (4 ug/mL) | 26,777 | 21,450 | 23,069 |
| Unstained, 33°C | 1,550 | 491 | 18,712 |
| 33°C | 4,101 | 1,666 | 20,220 |
| 33°C + Camptothecin (4 ug/mL) | 35,159 | 24,879 | 23,851 |

Supplemental Figure 6. Caspase 9 Flow Cytometry

(A-B) Flow cytometry histograms, median fluorescence intensities, median absolute deviations, and cell counts for activated caspase 9 were detected on a CytoFLEX LX cytometer with a fluorochrome inhibiter of caspases (FLICA) conjugated to a green FAM fluorescent probe. Camptothecin (4 ug/mL) was used for caspase 9 dependent apoptosis induction. Treatment conditions were (A) 10 μM Q-VD-Oph pan caspase inhibitor; (B) 10 mM galactose as the primary carbon source (initiated at least 3 days prior to and maintained through inoculation and incubation), 5 μM oligomycin, and 33ºC incubation temperature. Percent changes in median fluorescence intensities were presented in Figure 6.
